# Divergence in the Pelagic Zone: Genomic Signatures of Speciation and Adaptation in the Ctenophore *Mnemiopsis*

**DOI:** 10.1101/2024.10.10.617593

**Authors:** Remi N. Ketchum, Edward G. Smith, Leandra M. Toledo, Whitney B. Leach, Natalia E. Padillo-Anthemides, Andreas D. Baxevanis, Adam M. Reitzel, Joseph F. Ryan

## Abstract

**Background:** Understanding how populations diverge is one of the most compelling questions in evolutionary biology but our grasp on the genomic mechanisms underpinning divergence is limited to a handful of species. Indeed, we know even less about divergence in the pelagic zone, where barriers to gene flow are seemingly absent. The holopelagic ctenophore *Mnemiopsis leidyi* is the most widely used ctenophore in experimental biology and has become an important model system in studies ranging from developmental biology to neurobiology. In addition, its relatively small and tractable genome provides a powerful foundation for genomic and evolutionary analyses. However, we still lack a clear understanding of species boundaries, population structure, and the evolutionary forces shaping divergence within *Mnemiopsis*, limiting both evolutionary and ecological interpretations. To expand our general understanding of divergence across novel environments as well as resolve a long-standing taxonomic debate, we generated the most comprehensive genomic study to date of the holopelagic ctenophore *Mnemiopsis* across a large expanse of its native range.

**Results:** By leveraging multiple analytical approaches and generating two near-chromosome level genomes, we identify two distinct species of *Mnemiopsis* with high levels of genome-wide divergence along the US Atlantic coast, which correspond to *M. leidyi* and *M. gardeni*. Our demographic analyses suggest that *M. leidyi* and *M. gardeni* began to diverge during the mid-to-late Pleistocene climate transitions and were later shaped by post-glacial oceanographic changes. We highlight substantial genomic rearrangements and copy number variation between species as well as uncover key genes under selection that are likely important for environmental adaptation.

**Conclusions:** Together, these findings provide compelling evidence that the ctenophore currently recognized as *M. leidyi* represents more than one species. Recognizing cryptic species boundaries is critical for future study designs, environmental monitoring, and developing targeted management strategies. Altogether, we connect microevolutionary processes with macroevolutionary patterns and provide new insights into how ocean dynamics drive speciation and adaptation in pelagic ecosystems.

## Background

Speciation, the process through which populations diverge into groups that no longer exchange alleles [1], is a fundamental evolutionary process responsible for generating much of the biodiversity on earth. Understanding how populations diverge and eventually become reproductively isolated has been a central question in evolutionary biology for decades [2–4]. Two primary models have traditionally been invoked to explain this process. In the first, allopatric speciation, geographic barriers restrict gene flow, allowing populations to diverge independently. In the second, which includes parapatric and sympatric speciation, divergence occurs despite partial or complete geographic overlap [5, 6]. Although allopatric speciation is widely regarded as the predominant mode of speciation [3, 5, 7], its prevalence in marine environments is questionable given the scarcity of physical barriers and the high dispersal potential of many organisms, which can lead to patterns of extensive connectivity [8–11]. Indeed, understanding how species arise and become established is particularly challenging in marine environments, although many examples of coexisting marine sibling species suggest that marine speciation may often be sympatric. This process could be driven either by rapid adaptation to different ecological niches, genomic features that limit recombination and restrict introgression at specific loci (termed barrier loci; [12–15]), and/or hidden barriers to gene flow. While the development and spread of incompatible genotypes is central to both sympatric and allopatric speciation, the exact genomic regions and architecture which mediate this incompatibility, and the chronology of these genetic changes is still widely debated [11, 16].

The genomic patterns associated with speciation depend largely on whether divergence happens in the presence or absence of gene flow, either during the initial split or following secondary contact [12, 17, 18]. In allopatric speciation scenarios, divergence can proceed unfettered by the homogenizing effects of migration [18, 19]. Meanwhile, in speciation-with-gene-flow scenarios, divergence is constrained by the balance between the degree of genetic linkage and recombination versus the strength of selection or drift [17]. In this scenario, regions of low recombination (e.g., centromeric regions or structurally complex loci), can facilitate divergence by maintaining linkage among locally adapted alleles despite ongoing gene flow [20, 21]. In marine species where physical barriers to dispersal can be weak, genomic features which reduce recombination are likely strong drivers of genomic divergence. However, most studies to date have focused on single nucleotide polymorphisms (SNPs) and while informative, this has left the contribution of structural variants to speciation comparatively unexplored (although see [6, 20–23]). Structural variants (SVs – insertions, deletions, and other genomic changes affecting >50 bp) can occupy more of the genome than SNPs, have the potential to dramatically change phenotypes, and are often more likely to impact organismal fitness than SNPs because they are big enough to contain entire genes and other functional elements [21, 24]. SVs have been shown to lead to reproductive isolation through several mechanisms [25], including: underdominance of heterokaryotypes [26], suppression of recombination [27, 28], gene duplications causing intrinsic postzygotic isolation [29], and SVs acting as mutations of large effect [21]. Despite their potential to drive speciation, structural variants have remained understudied due to challenges in detecting and genotyping them with short-read sequencing [30]. Recent methodological advances, however, now enable the generation of chromosomal- or near-chromosomal-level genome assemblies, providing new opportunities to comprehensively investigate SVs and their evolutionary significance to speciation [12].

In addition to methodological biases, the vast majority of speciation literature is based on terrestrial species with small population sizes, clear physical barriers to gene flow, and internal fertilization, all traits which are expected to facilitate the emergence of reproductive isolation [31–33]. To fully understand the mechanisms driving speciation, we must broaden the range of studied organisms to include animals with diverse life history strategies, particularly pelagic invertebrates that spend their entire life in the water column and are characterized by high fecundity [34], large population sizes [35], and high dispersal potential. These traits are expected to produce weak genetic differentiation and promote panmixia [36] and indeed, this pattern is observed in many marine species [37–40]. Holopelagic species, such as *Mnemiopsis* (A. Agassiz, 1865), remain particularly understudied in this regard.

*Mnemiopsis* is a genus within the phylum Ctenophora with a native range that extends continuously along more than 25,000 km of the Atlantic coast of the Americas, from Kittery, Maine to the Valdes Peninsula in Argentina [41–43]. These comb jellies are considered to be one of the 100 most invasive species in the world [44] and their introductions into European waters have triggered community trophic cascades and collapse of fisheries [45]. The extensive geographic distribution and invasive abilities of *Mnemiopsis* are driven in part by its broad environmental tolerance [46], flexible planktivorous diet [47], extensive regeneration abilities [48], high fertility levels [49], and a capacity for self-fertilization [50]. Risk management for invasive species requires genetic information to establish management units and accurate taxonomic classification is a fundamental component of effective control measures. Further, many invasive species have the capacity for rapid adaptation [51] which, in turn, may make them more likely to speciate following a biological invasion although relatively few studies have examined this phenomenon [52].

Despite its rich 100-year history as an evo-devo model [53] and the availability of extensive genomics resources [54–57], our understanding of species boundaries in *Mnemiopsis* is subject to debate [58–62]. The World Register of Marine Species (WoRMS) currently divides *Mnemiopsis* into two accepted species: *Mnemiopsis leidyi* (A. Agassiz, 1865; type location Massachusetts) and *Mnemiopsis gardeni* (L. Agassiz, 1860; type location South Carolina). A third species, *Mnemiopsis mccradyi* (Mayer, 1900; type location South Carolina), is currently recognized as a synonym for *Mnemiopsis leidyi* [63]. However, there is much confusion as to the ranges and differences in morphology between these two named species which is further complicated by anthropogenic range expansion. Most modern literature considers all ctenophores along the Atlantic and Gulf coasts of the United States as a single species, specifically *Mnemiopsis leidyi* [64]. Earlier studies have raised the possibility of multiple *Mnemiopsis* species [58–62], but the molecular data available at the time were limited in either marker number or population sampling, making it difficult to resolve this question conclusively. With the compact genome of *Mnemiopsis* and the rapid democratization of next-generation sequencing (NGS) technologies, we can now examine population relationships at unprecedented resolution and address this long-standing controversy more comprehensively than was previously possible.

We conducted whole genome short read sequencing of 118 individuals from 13 populations within *Mnemiopsis’* native range along the United States Atlantic coast and generated two near-chromosomal length genome assemblies of a northern and southern representative of each genus. We applied a suite of genomic approaches to understand holopelagic population dynamics, demographic histories, changes in genomic architecture, and adaptation to new environments. Taken together, these findings highlight barriers to gene flow in holopelagic species, illuminate mechanisms that may underlie invasion success, and address a critical knowledge gap in marine speciation research, which has historically focused on a relatively narrow subset of species.

## Results and Discussion

### Whole-genome sequencing of Mnemiopsis populations

We collected and sequenced the genomes of 118 *Mnemiopsis* individuals from 13 populations from their native range along the US Atlantic coast (Figure 1A, Additional file 1: Table S1). All samples were collected from < 10m surface waters and preserved in ethanol within an hour of collection. We generated a total of 730 Gb of Illumina reads and an average genome depth of 18.89x. This sequence data was used to generate datasets for subsequent genomic analyses (see Methods).

**Fig. 1.**
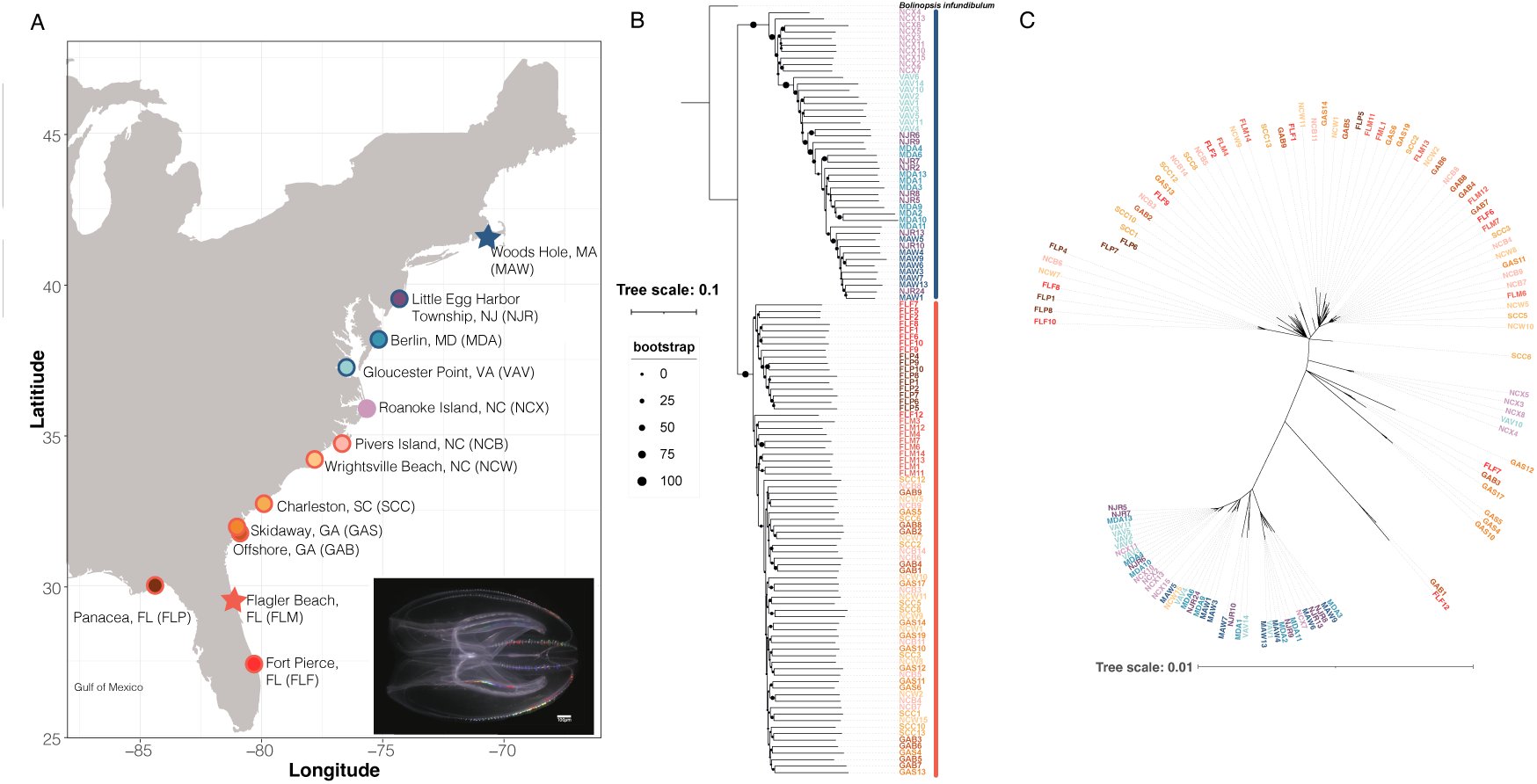
Mitonuclear discordance in *Mnemiopsis*. **A)** Sampling map showing all *Mnemiopsis* collection sites (site abbreviations in parentheses). The outer circle indicates the lineage assigned to each population (blue = northern; red = southern). Roanoke Island, NC is shown without an outer ring because it is later identified as a hybrid population. Stars denote populations also used for long-read PacBio HiFi genome sequencing and assembly. **B)** Nuclear phylogeny inferred from 1,302 SNPs using ngsDist and FastME, rooted with *Bolinopsis infundibulum*. Circle size at each branch reflects bootstrap support; both the northern and southern lineages showed 100% support. Sample ID colors correspond to sampling locations (matching panel A), and the colored bars to the right of each ID denote lineage membership (blue = northern; red = southern). **C)** Mitochondrial phylogeny generated from a whole-mitogenome alignment using IQ-TREE v2.3.2. Sample ID colors indicate lineage assignment.

We also produced two high-quality, near chromosome-level genome assemblies of one sample from Massachusetts (*M. leidyi*; northern lineage) and one sample from Florida (*M. gardeni*; southern lineage) using PacBio HiFi sequencing technology (Additional file 1: Table S2). When assembled, the *M. gardeni* genome was 206.6 Mb in length, with half of the genome assembled into contigs 3.6 Mb or larger (N50). The *M. gardeni* assembly was comprised of 174 contigs, with the 100 largest contigs making up 98% of the assembly. The *M. leidyi* genome was 215.8 Mb in length, with half of the genome assembled into contigs 2 Mb or larger. The *M. leidyi* assembly was comprised of 207 contigs, with the 100 largest contigs making up 89% of the assembly. We identified 27,570 and 26,046 protein-coding genes within the *M. gardeni* and *M. leidyi* genomes, respectively, and were able to functionally annotate 27,476 genes within the *M. gardeni* genome (Additional file 2: Table S3). Analyses using the Benchmarking Universal Single-Copy Orthologs program [65] revealed that our assemblies exhibit a similar degree of completeness to other ctenophore genomes ([54, 66], (Additional file 1: Table S2). We recovered 91% (83% complete and 8% partial) of the expected core genes for *M. leidyi* with 9% of them missing. For *M. gardeni*, we recovered 92% (84% complete and 8% partial) of the expected core genes, with 8% of them missing.

### Population genomic evidence of two Mnemiopsis species

We performed phylogenetic analyses using nuclear and mitochondrial markers from our *Mnemiopsis* samples collected along the Atlantic Coast. Our nuclear phylogenetic tree (Figure 1B) revealed the presence of two distinct clades that corresponded to the northern lineage (Woods Hole, MA to Roanoke Island, NC) and the southern lineage (Pivers Island, NC to Panacea, FL). We observed a similar two-clade pattern in our phylogenetic analysis of the complete mitochondrial genomes (published mitogenome is 10,326 bp [67]), driven largely by ∼100 nearly fixed nucleotide differences (Figure 1C). However, we identified six individuals from populations near the geographic interface of these two clades that exhibit mitonuclear discordance. Specifically, we observed one individual collected at Gloucester Point, Virginia (VAV10) and four individuals collected in Roanoke Island, North Carolina (NCX3–5,8) that had northern nuclear and southern mitochondrial genotypes. Likewise, we observed one individual collected from Wrightsville Beach, North Carolina (NCW15) that had a southern nuclear and northern mitochondrial genotype. This level of mitonuclear discordance could be consistent with hybridization in a secondary contact zone and early generation hybrids [62, 68].

To further investigate these admixture patterns, we evaluated the presence of 2–7 populations (by adjusting the K parameter) within our dataset using ngsADMIX [69]. Setting K=2 separates individuals into northern and southern lineages with the Roanoke Island, NC (NCX) and Gloucester Point, VA (VAV) samples showing signs of admixture (Figure 2A). When K=3, the Roanoke Island samples form their own subpopulation and, at higher K values, more substructure within the two lineages becomes evident (Additional file 1: Fig. S1). MDS plots (Figure 2B) support the results obtained using ngsADMIX, with the first principal component congruent with the patterns observed by setting K = 2 and accounting for 39.4% of the variance in the data. The average F_ST_ between the northern and southern lineage was 0.278 when Roanoke Island, NC was included in the northern lineage, and 0.315 when it was removed from the comparison. Our pairwise F_ST_ comparisons also highlight two distinct lineages (Additional file 1: Table S4). Due to these cumulative results, we also tested for evidence of hybridization in our data using *f*_3_ statistics [70]. We compared the Roanoke Island, NC (NCX) population to the most northern population (Woods Hole, MA; MAW) and most southern population (Panacea, FL; FLP) and, notably, this resulted in a negative *f*3 statistic and an absolute Z score >= 3 (threshold used for significance), highlighting a significant degree of admixture in the Roanoke Island, NC population (Additional file 1: Table S5). In line with a recent study which detected evidence of admixture in Virginia [62], we also observed signatures of admixture in Virginia (Gloucester Point, VA; VAV), as well as other northern populations (including in populations ∼450 km away, namely NJR and MDA). In contrast, there is no admixture observed in any populations south of Roanoke Island, even though Pivers Island, NC (NCB) is comparatively close (∼180 km south).

**Fig. 2:**
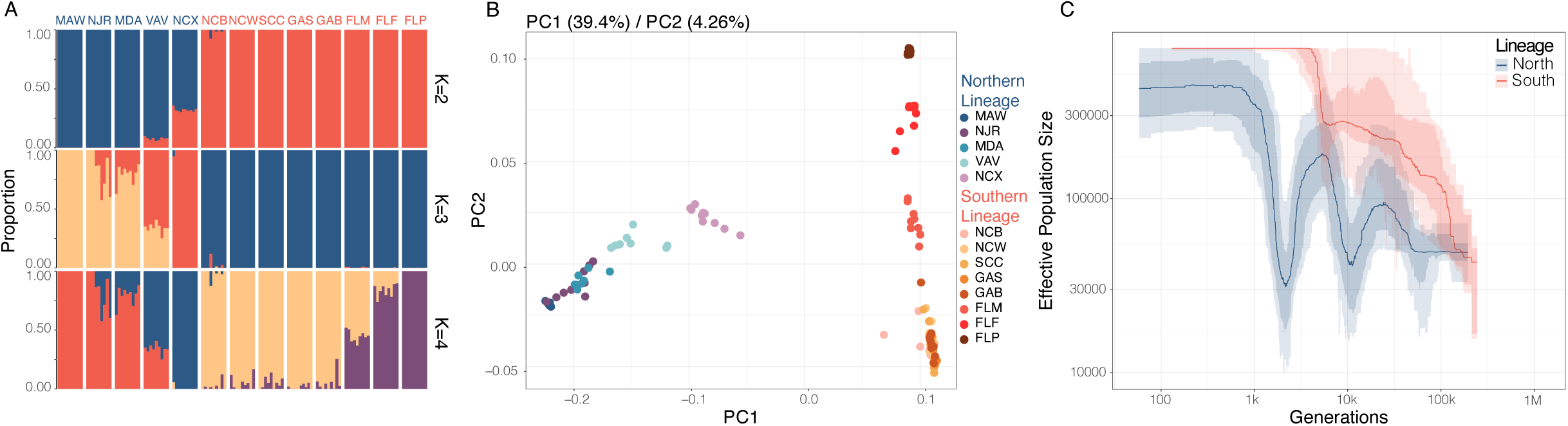
*Mnemiopsis* exhibits strong population structuring along the US Atlantic Coast. **A)** Admixture plots for the 118 *Mnemiopsis* samples for K = 2 to K = 4 where each bar represents a single individual, and the colors represent each of the K components. **B)** MDS plot for PC1 and PC2 with the percent of total variance explained by each component shown above the plot in parentheses. Samples are colored according to their sampling location with *M. leidyi* samples (northern lineage) clustering on the left and *M. gardeni* samples (southern lineage) clustering on the right. **C)** Stairway plot results show different demographic histories of the two species with a generation time of one year and a mutation rate of 6.85 × 10^-8^ per base pair per year.

Our analyses reveal two divergent genetic lineages in US Atlantic *Mnemiopsis*. These results are consistent with previous studies that reported genetic differentiation between Massachusetts and Florida populations [58] and identified a population breakpoint near Cape Hatteras, NC [59]. Building on this earlier work, our high-resolution spatial and genomic sampling further demonstrates that the genetic breakpoint between the two lineages is a narrow hybrid zone. There is cumulative evidence of admixture observed from Roanoke Island, NC up to Gloucester Point, VA (∼170 km north) but little evidence of admixture observed at Pivers Island, NC, which is located only ∼180 km to the south. Although *f*3-statistics suggest evidence of admixture in Little Egg Harbor Township, NJ (NJR), and Berlin, MD (MDA), this result contrasts with the other population genetic results and is therefore more likely explained by a counterintuitive property of *f-*statistic computations known as sister repulsion [71]. Sister repulsion can occur when there is unidirectional gene flow between study regions or when populations experience gene flow from an unsampled genetically distant source. Extending our approach to new geographic locations across the reported range of *Mnemiopsis* would make discerning between these two processes possible. However, we posit that this pattern is most likely explained by unidirectional gene flow given our demographic modeling results (see next section). Taken together, our findings indicate that the genetic breakpoint is located where the Gulf Stream leaves the coast and moves offshore into the open Atlantic Ocean, and we find evidence of a hybrid zone potentially resulting from recent secondary contact. We also see high levels of genome-wide divergence, which tends to be more typical of species that experienced geographic isolation during the early stages of speciation [72, 73], however demographic analyses are required to explore this further. With this in mind, we will hereafter refer to the northern lineage as *M. leidyi* and the southern lineage as *M. gardeni* (following the naming scheme employed in WoRMS).

### Demographic history of Mnemiopsis

We employed a Stairway Plot analysis [74] to reconstruct the demographic histories of *Mnemiopsis*, assuming a generation time of 1 year (see Methods). The two species show strikingly different population size trajectories beginning roughly 10,000 years ago (Figure 2C). Around this time frame, *M. gardeni* exhibits sustained population growth whereas *M. leidyi* displays a more complex history with two population bottlenecks, one around 10,000 years ago and a second between 2,000 - 3,000 years ago, followed by a recent recovery and expansion. The divergence between the effective population size curves for *M. gardeni* and *M. leidyi* that begins ∼10,000 years ago coincides with major post-glacial events. Around this time, the Laurentide Glacier retreated (∼8,000 years ago), and the sudden drainage of Lake Agassiz released a large influx of cold freshwater into the northern Atlantic [75]. This influx of cold freshwater likely reduced the current in the ocean conveyor belt and resulted in a decrease in the amount of warm seawater traveling northward, subsequently causing temperatures to drop in the Northern Hemisphere [75, 76]. The population expansion that followed may have been a result of a reduction in the Gulf Stream current following the Lake Agassiz flood which would have allowed *Mnemiopsis* to migrate northward. It is unclear what would have caused the population bottleneck ∼ 2,000 – 3,000 years ago in *M. leidyi*. It is possible that it was caused by the Little Ice Age – a cooling period in the northern hemisphere, although this occurred roughly between 1300 – 1850 [77, 78] and therefore does not line up perfectly with our estimated bottleneck.

Our results differ from the sequentially Markovian coalescent (PSMC) results reported in Jaspers et al. 2021 [58], which were based on a pairwise comparison of populations from Massachusetts and Florida. While both studies identify population expansions about ∼50,000 generations ago and a population bottleneck ∼10,000 generations ago, their *M. leidyi* samples did not show oscillating population size changes. Additionally, their *M. gardeni* samples showed a steady increase in population size from ∼500,000-50,000 generations ago and then a decrease in population size from then onwards. In contrast, ours has shown a steady increase in population size from 100,000 generations ago to present day. The difference in findings can likely be explained by the increased geospatial resolution in our study. In addition, while the PSMC method [79] performs well for inferring ancient histories, the Stairway Plot method is more accurate for inference of more recent population histories and we see the greatest discordance in our results when looking at more recent population histories [74, 80].

Our demographic analysis using *Moments* further refined our understanding [81]. The best-fit demographic model (sc3ielsm1, Additional file 1: Fig. S2-4) included secondary contact and three epochs in each species following their initial divergence. The split between *M. leidyi* and *M. gardeni* occurred approximately 223 Kya, followed by ∼107 Kya of low-to-moderate bidirectional gene flow. The second epoch began around 116 Kya and was characterized by genetic isolation lasting roughly 89 Kya. The most recent epoch started approximately 27 Kya, during which asymmetrical migration occurred predominantly from *M. gardeni* to *M. leidyi*. Across these epochs, *M. leidyi* experienced an initial population expansion, followed by a sustained decline, whereas *M. gardeni* displayed the opposite pattern, an initial contraction followed by consistent population growth.

Together, Stairway Plot and *Moments* analyses reveal broadly consistent demographic patterns. *Moments* infers a reasonably ancient divergence time, perhaps because incorporating migration into the model allows some population differences to be explained by a more ancient split rather than recent divergence. Both approaches recover evidence of pronounced population size reductions in *M. leidyi*. The magnitude and number of contractions differ between methods, likely because *Moments* is limited to modeling a maximum of three population size changes, whereas Stairway Plot can recover more complex histories. These methods also concur that *M. gardeni* has undergone a recent expansion, though they differ on whether *M. leidyi* has rebounded or continues to experience a population contraction. Overall, the demographic models suggest that divergence likely began in the mid-to-late Pleistocene, potentially caused by distant colonization events, geological change, sea level changes or changes in prevailing currents [82]. However, it remains unclear whether genetic isolation was initiated by glacially driven flooding and subsequent re-strengthening of the Gulf Stream, or whether divergence predates these events, with post-glacial oceanographic changes instead facilitating secondary contact and asymmetric migration from *M. gardeni* to *M. leidyi*.

### Structural genomic evidence of speciation

Chromosome-scale linkages of orthologous genes tend to be conserved even over very long time frames [83–87]. As such, chromosomal rearrangements are major evolutionary events that can either cause speciation events and/or provide evidence of long periods of isolation between lineages. To investigate structural differences between *M. leidyi* and *M. gardeni* genomes, we produced two high-quality, near chromosome-level genome assemblies of one *M. leidyi* sample from Massachusetts and one *M. gardeni* sample from Florida. We conducted macrosynteny analyses to check for evidence of chromosomal rearrangements between the two species’ near-chromosomal length genome assemblies. In addition, we compared both genomes to the recently published chromosome-level genome assembly of *Bolinopsis microptera* [86], a lineage that is closely related to *Mnemiopsis* [88], in order to infer ancestral chromosomal states. We identified at least one chromosomal rearrangement between the *M. gardeni* and *M. leidyi* genomes (Figure 3A-C). The *M. leidyi* scaffold L006 maps to two *M. gardeni* scaffolds: G002, which maps completely to chromosome one of *B. microptera*, and G004, which maps completely to chromosome eight of *B. microptera*. The fact that this linkage group is present in both *M. gardeni* and *B. microptera* suggests that the genomic rearrangement occurred in *M. leidyi*. There are potentially other rearrangements involving the *M. leidyi* scaffolds L012 and L018, but a chromosome-scale assembly is needed to confirm or refute those possibilities.

**Fig. 3:**
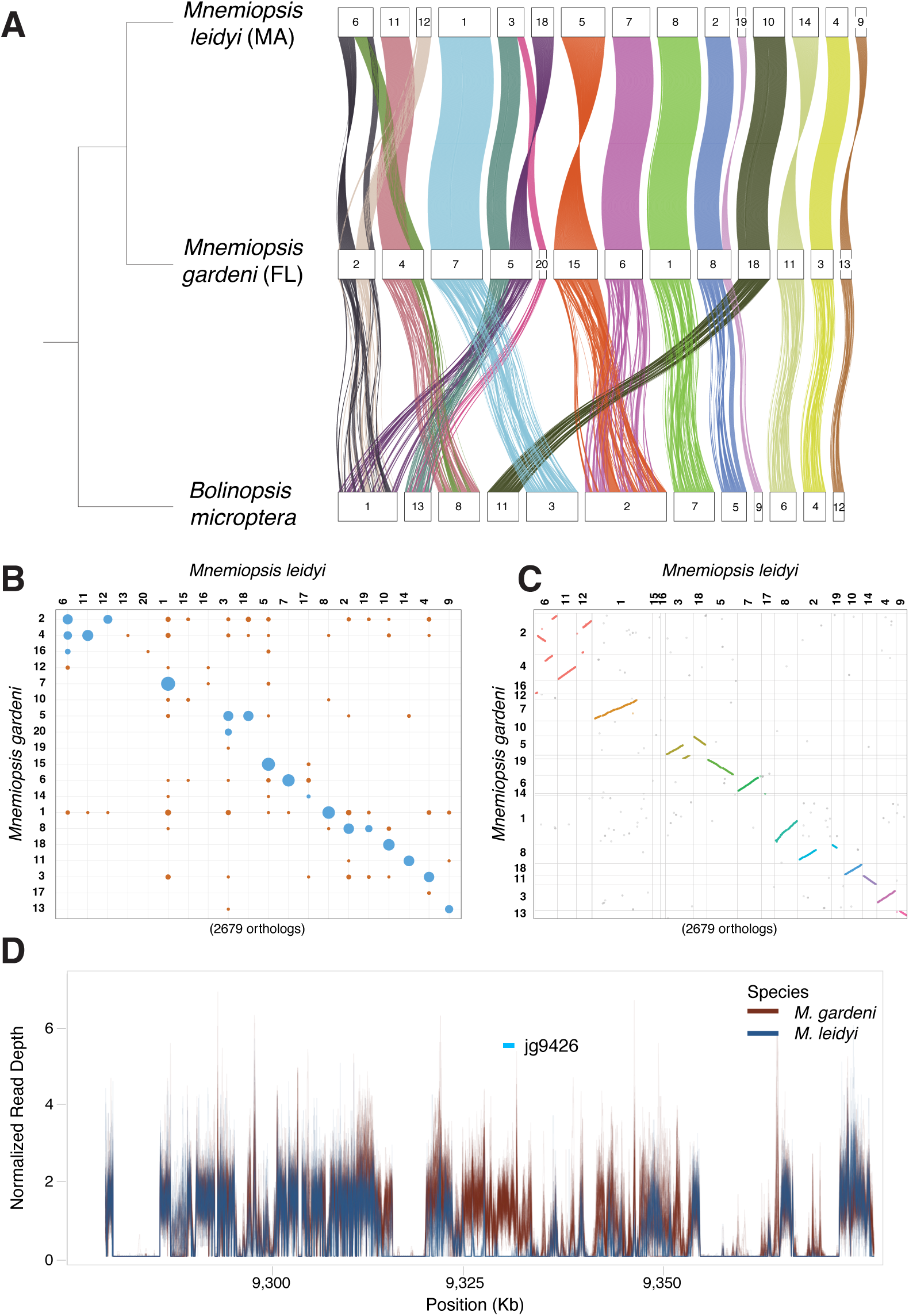
Large-scale genomic rearrangements between species. **(A)** Chord diagram of macrosynteny on a near-chromosome scale between *Mnemiopsis leidyi* (top), *M. gardeni* (middle), and *Bolinopsis microptera* (bottom). The coordinates of 12,134 orthologs (based on reciprocal best BLAST hits) are plotted. **(B)** Plot of macrosynteny between *M. leidyi* and *M. gardeni*. **(C)** Oxford dot plot between *M. leidyi* and *M. gardeni*. Each dot represents an ortholog. **(D)** The genomic landscape neighboring the top copy number variation outlier gene based on fixation index values (*V*_ST_). Normalized read depth plot for short-read sequences from *M. leidyi* (blue) and *M. gardeni* (red) for *M. gardeni* scaffold 001 from position 9,278,170 – 9,378,170.

As copy number variants (CNVs) can drive genome evolution [89] and contribute to both speciation events [90] and environmental adaptation [91], we examined CNV patterns between the two species. Genome-wide patterns of CNV across samples largely mirror that of SNP variation, with separation of samples into clusters associated with *M. leidyi*, samples south of the Outer Banks, and the Floridian samples (Additional file 1: Fig. S5). We assessed the 67 genes in the top 1% of fixation index values (*V*_ST_; Additional file 3: Table S6). Of these 67 genes, only 21 (31%) were assigned a gene ontology (GO) category [92] based on sequence similarity (Additional file 1: Table S7). We ran a Monte Carlo analysis and showed that this is significantly less (*P* = 0.0006) than would be expected by chance. On average, a random set of 67 genes would produce 32 (48%) genes within an associated GO category. These data suggest that the CNV outliers are enriched in so-called “orphan” genes. This is not unexpected, as “orphan” genes are often associated with effector functions (e.g., [93]) rather than highly conserved functions that are often critical to survival.

The top *V*_ST_ CNV gene (jg9426 in the *M. gardeni* genome) is a putative G-protein coupled receptor (GPCR), a class of cell surface receptors that detect molecules (e.g., light energy, lipids, proteins) and activate cellular responses. This GPCR gene copy is missing from *M. leidyi* and appears to be the result of either a species-specific duplication in *M. gardeni* or from a deletion of an ancestrally duplicated gene in *M. leidyi* (*M. gardeni* gene IDs: jg9426 and jg9432, *M. leidyi* gene ID: ML07987a; Figure 3D, Additional file 1: Fig. S6). It is one of four predicted GPCRs in these 67 genes (Additional file 1: Table S8). These GPCRs are all part of GPCR clades that have independently radiated in ctenophores and have a many-to-many relationship with non-ctenophore GPCRs, making it difficult to infer function. Loss or duplications in these genes would likely lead to changes in sensory capabilities or cellular responses downstream of sensory inputs, changes that are widely hypothesized to facilitate ecological speciation (speciation caused by divergent selection; [94, 95]). These microevolutionary changes are consistent with the macroevolutionary pattern of loss and expansion observed in GPCRs, offering insight into mechanisms that underly the independent radiation commonly seen in phylogenetic analyses of channel and receptor genes from multiple species. These observations are consistent with the “accordion model” of gene evolution, where rounds of extensive loss are followed by rounds of extensive duplication [96]. Overall, patterns in the CNV data mirror those seen at the single nucleotide level and, although gene loss in *M. leidyi* may be a result of severe bottlenecks or selective pressures, it may also stem from reference bias, highlighting the need for pangenomic approaches [97, 98].

### Genomic signatures of selection in genes mediating environmental sensing

To investigate signatures of selection and local adaptation, we used the cross-population composite-likelihood ratio (XP-CLR) test [99]. We observed key sensory related and neural genes found in regions within the top 10% of XP-CLR values (Additional file 4: Table S9). These included 79 GPCR’s, four TRP channels, five glutamate receptors, the single *Mnemiopsis* voltage-gated calcium channel, one of the two *Mnemiopsis* voltage-gated sodium channels, 10 voltage-gated potassium channels, three acid-sensing ion channels (ASIC), two calcium-activated potassium channels, a voltage-gated chloride channel, and a DEG/ENaC channel (Additional file 1: Tables S10-11). Our outlier dataset also contains many other genes that are relevant for nutrient sensing and metabolism; these include the mechanistic target of rapamycin (mTOR), acetyl coenzyme A synthase, malate dehydrogenase, and cytochrome c oxidase subunit 2 genes.

The top XP-CLR outlier region contains the *syntaxin-6* gene (jg7623), which promotes the movement of transport vesicles to target membranes. We also observe the *O-GLcNAcase* (jg8681) gene in the outlier dataset, which codes for a post-translational modifier of serine and threonine residues on a wide range of proteins [100]. We highlight these two genes in Figure 4 as they have both been implicated in modulating autophagic flux in response to nutrient status in *Caenorhabditis elegans* [101, 102]. Both sweep regions show multiple signatures of selection, including reduced nucleotide diversity in *M. leidyi*, elevated F_ST_ values, and an excess of rare alleles (as evidenced by the more negative Tajima’s *D* values for *M. leidyi*). We hypothesize that overwintering in northern populations may be a driver of these selective sweeps, as organisms are likely nutrient-deficient during this time. Interestingly, *M. gardeni* populations have significantly lower Tajima’s *D* values than *M. leidyi* populations across the entire genome (W=1.16 × 10⁹; *P* < 0.01). However, this pattern breaks down at the target of selective sweeps. This may be consistent with overall genomic patterns highlighting a population expansion in *M. gardeni* and population contractions in *M. leidyi* that is disrupted at the site of selective sweeps.

**Fig. 4:**
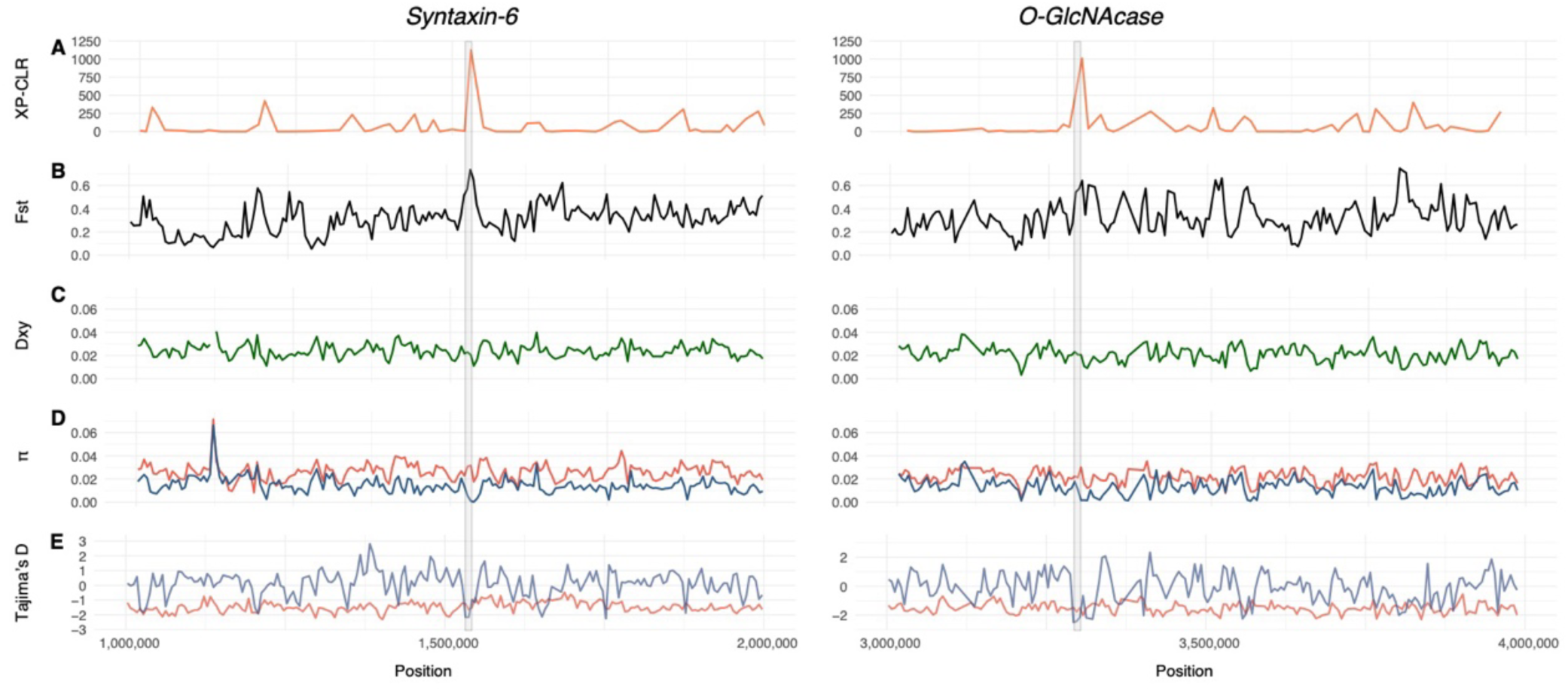
Genes involved in autophagic flux are under strong positive selection between species. The two genes: *syntaxin-6* (left) and *O-GlcNAcase* (right) that are under putative positive selection. The (A) XP-CLR, (B) F_ST_, (C) D_XY,_ (D) nucleotide diversity (π), and (E) Tajima’s *D* values were calculated with nonoverlapping windows and step sizes. The locations of the two sweeping windows are shown with gray boxes. For panels D and E, blue lines indicate *Mnemiopsis leidyi* and red lines indicate *M. gardeni*.

To highlight functional categories of these genes, we conducted gene ontology (GO) analysis [92] of the 2,093 genes that overlap with the genomic windows ranked in the top 10% of XP-CLR values (Additional file 1: Table S12). These genes were significantly enriched (*P* <= 0.05) for GO molecular function categories, including those associated with sensory processing, neural activity, or diet, such as GPCR signaling pathways (*P* = 0.0136), GPCR activity (*P* = 0.0152), Golgi vesicle transport (*P* = 0.0399), neurotransmitter transport (*P* = 0.0424), and chitin metabolic processes (*P* = 0.0056). The chitin metabolic processes category includes two chitinase genes, which are critical for digesting prey. These findings suggest that recent selection has acted on genes involved in environmental sensing and resilience to nutrient deficiency and provide important candidate genes for future functional verification.

## Conclusions

Both the population genomic and structural genomic evidence are consistent with there being two species of *Mnemiopsis* on the US Atlantic coast. We report high levels of genome-wide divergence, as well as mitonuclear discordance. Demographic modeling suggests that these species diverged in the mid-to-late Pleistocene, with long-term genetic isolation followed by secondary contact and asymmetric migration from *M. gardeni* into *M. leidyi*, a scenario supported by the hybrid zone we identify along the North Carolina coast. We highlight large-scale genomic rearrangements between species that are consistent with a history of population contractions or small founder populations. We find evidence of selection on sensory and nervous system genes, with similar trends being reported in other species following speciation events [88, 103–106].

Further research is needed to investigate morphological, reproductive, and life-history traits that may help provide further evidence for multiple species, as well as to examine the possibility that more *Mnemiopsis* species exist in Central and South America, as the mitochondrial results of Bayha et al. suggest [59]. More broadly, our knowledge of evolution is heavily biased across the tree of life, and this study reinforces the importance of expanding taxon sampling as a means towards gaining a much richer understanding of how evolutionary forces shape the genomic variation between populations and species.

## Methods

### Sampling

One individual was collected from each of two locations for genome assembly: Woods Hole, MA and Flagler Beach, FL. These samples were collected in August 2023 from surface waters (<2m) using a cteno-dipper [49]. Samples were preserved in ethanol and placed in a −20°C freezer until further processing.

A total of 118 *Mnemiopsis* samples were collected from thirteen locations along the Atlantic and Gulf coast of the United States. These locations included: Woods Hole, MA, Little Egg Harbor Township, NJ, Berlin, MD, Gloucester Point VA, Roanoke Island, NC, Pivers Island, NC, Wrightsville Beach, NC, Charleston, SC, Skidaway Island, GA, Skidaway Island offshore, GA, Flagler Beach, FL, Fort Pierce, FL, and Panacea, FL (Figure 1). Samples were collected between August 2021 and November 2022 from surface waters (<10m) using either a cteno-dipper or a tow net. Samples were immediately preserved in ethanol and placed in a −20°C freezer until further processing.

### DNA Extraction and Sequencing

High molecular weight DNA was extracted according to the CTAB/chloroform protocol detailed in Erlich et al. 2017 [107]. DNA extractions were visualized on a 1% agarose gel for quality control and quantified using a Qubit dsDNA High Sensitivity Assay Kit on a Qubit 2.0 Fluorometer. For the genome assemblies, sequencing was performed on a Pacific Biosciences Sequel IIe platform, in the Circular Consensus sequencing (CCS) mode. A total of 17.7 - 20.2 Gb of HiFi reads were generated using CCS for the two individuals. PacBio library preparation and sequencing were performed at NIH Intramural Sequence Center (NISC) in Rockville, MD.

For short-read whole genome sequencing, all DNA extractions were visualized on a 1% agarose gel for quality control. One hundred and eighteen individually indexed, paired-end libraries with an approximate insert size of 550 bp were constructed using the NEBNext Ultra II FS DNA Library Preparation kit. Whole-genome sequencing was performed at the University of Florida’s ICBR Bioinformatics Core Facility on a single flowcell lane of a NovaSeq6000.

### De novo Genome Assembly

Hifiasm v0.19.8 [108] was used to assemble the two genomes. For the Flagler Beach, FL individual, the `--hom-cov` flag was set to 90. To reduce the amount of duplication, PurgeHaplotigs v1.1.2 [109] was run with the ‘contigcov’ option for low coverage, low point between the peaks, and high coverage cutoffs set to 10, 70, and 175 respectively. The percent cutoff for identifying a contig as a haplotig (`-a` flag) was set to 50% and this assembly was subsequently run through PurgeHaplotigs ‘clip’ option that identifies and trims overlapping contig ends. For the Woods Hole, MA individual, the `--hom-cov` flag was set to 101, the ‘contigcov’ options were set to 25, 70, and 155, and the `-j` and `-s` flags were both set to 55. The percent cutoff `-a` flag was set to 60% and the final assembly was also clipped to remove overlapping contig ends.

### Gene Prediction and Annotation

Genomic repetitive elements were identified with RepeatModeler v1.0.11 [110] to generate a *Mnemiopsis*-specific repeat element library. Repetitive regions were soft masked prior to gene prediction and annotation using RepeatMasker v4.0.8. We aligned previously published bulk RNA-Seq reads from tissues (PRJEB28334; [111]) to the above genome assembly using Bowtie2 [112]. Aligned RNA-seq data and gene models for *M. leidyi* (ML2.2.aa) were input into Braker v2.1.5 [113]. Assembled gene/protein models were functionally annotated using BLASTP v2.9.0+ with three protein databases: UniProt Knowledgebase Swiss-Prot protein models v2021-03, RefSeq invertebrate protein models, and the *M. leidyi* protein models [54]. Finally, BUSCO v5 [65] was used to measure the completeness of the genome assembly and protein sequences using the Eukaryota database.

### Macrosynteny Analysis

We analyzed conserved chromosome-scale synteny (macrosynteny) between these new Florida and Massachusetts *Mnemiopsis* assemblies, as well as the published *Bolinopsis microptera* genome assembly. To identify conserved genes, we performed diamond BLAST [114] to compute the reciprocal best BLAST hits between the 2013 ML2.2 *Mnemiopsis* gene models and the gene models of each of the new *Mnemiopsis* assemblies and the published *B. microptera* gene models. These reciprocal best BLAST hits were used to identify significantly conserved macrosyntenic blocks in MacrosyntR (version 0.3.3; [115]).

### Copy Number Analysis

Copy number variation between the two species was assessed using the GATK germline CNV (gCNV) pipeline in cohort mode [116]. The pipeline was run using the exome approach to focus specifically on genic regions and gene intervals were padded by 250 bp. Intervals with fewer than 50 reads in greater than 50% of individuals were excluded and samples from Roanoke Island in North Carolina were excluded due to their admixture. Denoised copy number ratios were processed in R to remove copy number invariant loci, perform principal component analysis, and to calculate the *V*_ST_ statistic. Loci with the top 1% of *V*_ST_ values were considered outliers. Microsynteny plots for CNV loci were generated by performing contig alignments in MUMmer [117]. SYRI was subsequently used to identify synteny and structural rearrangements, with plots generated using Plotsr [118, 119] and read depth was calculated using Samtools v.1.9 [120].

### Data Processing and Filtering of Short-Read Data

Raw sequence data was trimmed using Trimmomatic v0.39 [121] with a minimum phred score of 33. Once the data was filtered and adaptors were removed, the paired sequences were aligned to the Flagler Beach, FL genome using BWA-mem v0.7.12 [79]. Sequence alignments were filtered using Samtools for a minimum quality score of 30, Sambamba v0.6.6 [122] was used to filter duplicate reads with default parameters, and bamutil v1.0.15 [123] was used to clip overlapping read pairs.

We generated three datasets for our different analyses: LD-Filtered (89,750 SNPs), Lineage-Specific Filtered (3.8 million SNPs), and the All-Sites Filtered (81 million positions). For each dataset, we used the Flagler Beach, FL genome as our reference and ancestral genome as it was the most contiguous assembly.

#### LD-Filtered Dataset

This dataset was produced by using ANGSD v0.921 [124] to call SNPs and estimate genotype likelihoods using the Samtools model `(-GL 1`). We filtered the SNP data for 1) positions that were present in at least 94 out of 118 samples, 2) a maximum coverage of 2,360x per site across all individuals to avoid calling sites in highly repetitive regions, 3) a minimum of 354x coverage per site across all individuals, 4) a minimum base quality score >20, 5) a *P-*value cut off of 10^-6^ for calling polymorphic loci and 6) retained only SNPs with a global minor allele frequency >5%. The ANGSD output was then subsampled for one SNP out of every 50 and pairwise linkage disequilibrium (LD) was estimated with ngsLD v1.1.1 [125] on this subsampled dataset. Linked sites were pruned with the script ‘prune_graph.pl’ with a maximum distance of 10 kb between SNPs and a minimum weight of 0.5 [126]. The final LD-Filtered dataset was used to generate phylogenetic trees, ngsADMIX and MDS plots, and to calculate population pairwise F_ST_ statistics.

#### Lineage-Specific Filtered Dataset

The second dataset was filtered using the same ANGSD flags as the first dataset (excluding the LD filtering steps) but then underwent a second round of lineage-level filtering. Specifically, sampling locations from Woods Hole, MA to Roanoke Island, NC were included in the ‘northern lineage’ group and sampling locations from Pivers Island, NC to Panacea, FL were included in the ‘southern lineage’ group. These two groups were filtered separately and then later recombined. For the northern lineage, we filtered the dataset for 1) positions that were present in at least 36 out of 45 individuals, 2) a maximum coverage of 900x coverage per site across all individuals, 3) a minimum of 108x coverage per site across all individuals. For the southern lineage we filtered the dataset for 1) positions that were present in at least 58 out of 73 individuals, 2) a maximum coverage of 1,460x coverage per site across all individuals, 3) a minimum of 175x coverage per site across all individuals. We restricted the secondary lineage-specific filtering step to the subset of positions that were retained in the first round of filtering using the sites option in ANGSD. Filtering in this manner guarantees that sites with an allele that is fixed in one lineage will be retained. The final lineage-specific filtered dataset was used to generate sliding window F_ST_ values, XP-CLR values, and to calculate *f*_3_ statistics.

#### All-Sites Filtered Dataset

The third dataset was filtered with the same method as the Lineage-Specific Filtered dataset, but we included all non-polymorphic sites. This dataset was used as input to our demographic analyses and to calculate nucleotide diversity [127], Tajima’s *D* [128] and D_XY_ values.

### Population Structure Analysis

We used the ngsDist option in ngsTools [129] on the LD-Filtered dataset to compute genetic distances between individuals and perform multiple dimensional scaling (MDS). We then looked for signatures of admixture using ngsADMIX [69] with runs ranging from K = 2 to K = 7. Weighted pairwise F_ST_ values were estimated between the individuals from the North (Woods Hole, MA to Roanoke Island, NC) and the individuals from the South (Pivers Island, NC to Panacea, FL) using ANGSD with the `-doSaf 1` flag. We then used realSFS to estimate the 2D site frequency spectrum for each population and calculated the average pairwise weighted F_ST_ using `realSFS fst`. Weighted pairwise F_ST_ estimates were also calculated between each sampling location using ANGSD with the same parameters.

NgsDist was used to compute genetic distances from genotype probabilities with *Bolinopsis infundibulum* as the outgroup (PRJNA818620; [86]). Raw *Bolinopsis* sequences were filtered according to the same protocol as was applied to *Mnemiopsis* and then we ran ANGSD with *Bolinopsis* included, a minor allele frequency filter of 0.05 and a SNP *P*-value cut off of 1×10^-6^. This set of filters resulted in 1,302 SNPs that were used to generate the phylogenetic tree based on the nuclear genome. NgsDist was run with 100 bootstrap replicates and a boot block size of 20. Finally, FastME v2.1.6.1 [130] was used to visualize the pairwise genetic distances and RAxML-NG v0.9.0 [131] was used to place bootstrap supports onto the tree. Final figures were generated using the Interactive Tree of Life tool v6.9.1 [132].

Novoplasty v4.3.1 [133] was used to extract the mitochondrial genome from each trimmed sample using the published *Mnemiopsis leidyi* COI sequence (KF435121.1 [134]) as the seed. When this was unsuccessful, we instead used the published *Bolinopsis microptera* (PRJNA716277 [88]) whole genome sequences and ran Novoplasty with the published *B. microptera* COI seed (MW735734.1 [135]) to extract the complete mitochondrial genome. The complete *B. microptera* mitochondrial genome was then used as a seed for the remaining samples that initially failed Novoplasty. Using this approach, we were able to successfully assemble 112 out of 118 complete *Mnemiopsis* mitochondrial genomes. Next, we used MARS [136] to ensure that each mitochondrial genome began at the same location using the branch and bound method and the block length set to 100. We used MAFFT v7.480 [137] to generate an alignment and IQ-TREE v2.3.2 [138] to generate a phylogenetic tree.

### Demographic Analysis

To infer the demographic history of these species, we used two approaches based on allele frequency spectra. First, StairwayPlot v2 [74] was used to reconstruct effective population size estimates based on the folded SFS generated from the All-Sites Filtered dataset. Stairway Plot is an unsupervised analysis method that does not require prespecified demographic models. We used a mutation rate of 6.85 × 10^-8^ per base pair per year [58] and a generation time of 1 year [139]. The hermaphroditic *Mnemiopsis* can reproduce within the first month of its lifespan [140], its lifespan is approximately 2 years [139], and there is no information indicating that they ever lose the ability to reproduce. As such, the mean age of reproduction was set to 1 year but due to the surrounding uncertainty, we plotted the results in generations.

Second, we used the *Moments* Python library [81] based on the two-dimensional AFS to estimate population size changes and the timing and magnitude of introgression between each pair of genetic lineages. We subsampled our data so that we only included four populations, two *M. leidyi* populations (Woods Hole, MA and Little Egg Harbor, NJ) and two *M. gardeni* populations (Fort Pierce, FL and Panacea, FL), to conservatively remove putative hybrids and for computational efficiency. This dataset returned approximately 21 million sites in total. Unlike the Stairway Plot analysis, the *Moments* workflow requires demographic models to be specified at the start of the analysis. Here, we explicitly tested 102 demographic models (described in detail at https://github.com/z0on/AFS-analysis-with-moments/) that represent a wide range of possible demographic histories. Following the *Moments* pipeline described in Rippe et al., 2021, we first generated 100 bootstrapped two-population AFS datasets [141]. For a subset of 10 of these bootstrapped AFS, each of the 102 models was run six times using randomized starting parameters. Model log likelihoods were converted to Akaike’s information criterion (AIC) which were then used to find the best-fit run out of the six runs for each bootstrapped replicate. From the resulting set of AIC values, the demographic model with the lowest median AIC value across all 10 bootstrapped AFS replicates was chosen. Finally, the ‘winning’ demographic model was fitted to all 100 bootstrapped AFS to evaluate parameter uncertainties. As with the Stairway plot analysis, we assumed a mutation rate of 6.85 × 10^-8^ per base pair per year and a generation time of 1 year.

### F_3_ Statistics

We used *f*_3_ statistics, included as part of the ADMIXTOOLS package v.7.0.2 [70], as a test for detecting hybridization. The *f*_3_ statistic is a tool to test whether a population is admixed between two source populations or to measure shared genetic drift between two test populations compared to an outgroup population. If the *f*_3_ statistic is negative, this suggests that admixture has likely occurred. For this analysis, all populations which belonged to the northern mitochondrial lineage were combined into one population and all the populations which belonged to the southern mitochondrial lineage were combined into the second population. These two lineages were tested against Roanoke Island, NC and Gloucester Point, VA.

### Genomic Signatures of Selection

To detect regions of the genome under putative selection between the two lineages, we used XP-CLR [99] which uses the AFS to calculate local deviations between populations. We used a sliding window and step size of 10 kb with the `maxSNPs` flag set to 200. The genomic regions in the top 10% of the XP-CLR values were considered to represent selective sweeps and were associated with overlapping genes using Bedtools v.2.29.0 [142]. We used InterProScan (version 5.25-64) to associate gene ontology categories with the *M. gardeni* gene models. We conducted a Monte Carlo analysis to identify overrepresented gene ontology terms as was conducted in Babonis et al. 2018 [111].

To further characterize genomic patterns of regions under selection, we generated sliding window scans for F_ST_, D_XY_, nucleotide diversity (π), and Tajima’s *D* values, with the hybrid Roanoke Island, NC population removed. The Lineage-Specific Filtered dataset was used to generate the F_ST_ values. We used `realSFS F`_ST_ `index` in ANGSD to compute per-site F_ST_ indexes and `realSFS F`_ST_ `stats2` to perform a sliding window analysis with the window and step size set to 5 kb to ensure nonoverlapping windows. The All-Sites Filtered dataset was used to generate nucleotide diversity, Tajima’s *D,* and D_XY_ values. For nucleotide diversity and Tajima’s *D* calculations, we used `realSFS saf2theta` in ANGSD to generate folded SFS’s and then calculated these values along a sliding window (window size 5 kb, step size 5 kb) with `thetaStat do_stat`. For D_XY_ calculations, we used the CalcDxy.R code by Joshua Penalba in ngsTools [129] and then averaged the resulting values over the same 5 kb windows as the F_ST_ and nucleotide calculations. By combining several tests and summary statistics, we can make inferences about evolutionary forces acting on regions of interest with greater certainty.

## Abbreviations

SNP: Single Nucleotide Polymorphism
SV: Structural Variant
MDS: Multidimensional Scaling
PSMC: Pairwise Sequentially Markovian Coalescent
AIC: Akaike’s Information Criterion
GO: Gene Ontology
COI: Mitochondrial Cytochrome C Oxidase subunit I
LD: Linkage Disequilibrium
CNV: Copy Number Variation

## Declarations

## Acknowledgements

We would like to thank Dr. Ann Tarrant, Roland Hagan, Robert Aguilar, Dr. Deborah Steinberg, Tor Mowatt-Larssen, Eli at Outer Banks Kayaking Adventures, Ken Song, Carol Smith, Dr. Joshua Osterberg, Dr. Jacob Warner, Dr. Daniel Sasson, Dr. Marc Frischer, Dr. Adam Greer, Kyle Aaron, Dr. Holly Sweat, and Dr. Yuriy Bobkov for help collecting *Mnemiopsis* samples along the US Atlantic coast. We also thank Dr. Dan Schrider and Austin Daigle at UNC Chapel Hill for help analyzing and interpreting demographic modeling results. We would like to acknowledge the ICBR for their excellent sequencing and service (RRID:SCR_019152). We would like to thank Dr. James Thomas at the NIH Intramural Sequencing Center (NISC) for his help with the PacBio sequencing. We would also like to thank Dr. Gustav Paulay and Dr. Claudia Mills for their help navigating the complex taxonomy of *Mnemiopsis*. Finally, we want to thank Dr. Mark Ravinet for his help brainstorming ideas pertaining to the hybridization of these two species.

## Consent for Publication

The authors have all been consulted on draft and final versions of the manuscript and have consented to publication.

## Data Availability

The datasets supporting the conclusions of this article are available at the NCBI under BioProject accession number PRJEB87530. The genome assemblies for each *Mnemiopsis* species are also available under the above BioProject accession codes. All scripts are publicly available at https://github.com/remiketchum/Mnemiopsis_2026.

## Authors’ Contributions

Conceptualization: R.N.K and J.F.R. Investigation (fieldwork): R.N.K, W.B.L, E.G.S, J.F.R. Investigation (analysis): R.N.K, E.G.S, J.F.R, L.M.T, N.E.P. Resources: J.F.R, A.D.B, A.M.R. All authors read and approved the final manuscript.

## Ethics Approval and Consent to Participate

Not applicable.

