## Supplementary Materials for "Divergence in the Pelagic Zone: Genomic Signatures of Speciation and Adaptation in the Ctenophore *Mnemiopsis*"

**Supplemental Table 1.** Collection information for all samples collected along the US Atlantic coast including date of collection, number of samples collected, coordinates, and collection method.

|  | Date of Collection (yyyy-mm-dd) | # of samples collected | Coordinates | Collection Method |
| --- | --- | --- | --- | --- |
| Woods Hole, MA | 2021-08-16 | 15 | 41.524844, -70.674237 | Cteno-dipper (surface) |
| Little Egg Harbor Township, NJ | 2021-08-19 | 15 | 39.520152, -74.318928 | Plankton net (surface) |
| Berlin, MD | 2021-08-22 | 13 | 38.211082, -75.166615 | Cteno-dipper (surface) |
| Gloucester Point, VA | 2022-01-26 | 17 | 37.247772, -76.499840 | Cteno-dipper (surface) |
| Roanoke Island, NC | 2022-08-27 | 16 | 35.902007, -75.669288 | Cteno-dipper (surface) |
| Pivers Island, NC | 2022-11-15 | 16 | 34.717565, -76.670585 | Cteno-dipper (surface) |
| Wrightsville Beach, NC | 2022-08-31 | 15 | 34.208403, -77.796896 | Cteno-dipper (surface) |
| Charleston, SC | 2021-08-28 | 15 | 32.752802, -79.898984 | Cteno-dipper (surface) |
| Skidaway, GA | 2022-07-08 | 17 | 31.989390, -81.024077 | Cteno-dipper (surface) |
| Skidaway, GA (offshore) | 2022-09-19 | 9 | 31.924105, -80.966833 | <10m tow |
| Flagler Beach, FL | 2022-11-01 | 15 | 29.516855, -81.146889 | Cteno-dipper (surface) |
| Fort Pierce, FL | 2022-11-02 | 15 | 27.450034, -80.321657 | Cteno-dipper (surface) |
| Panacea, FL | 2022-03-29 | 10 | 30.023938, -84.385514 | Gulf Specimens |

**Supplemental Table 2:** Comparison of genome assembly benchmarks between the two published *Mnemiopsis leidyi* genomes and the two new genome assemblies.

|  | *Mnemiopsis leidyi* MA Ryan et al 2013 | *Mnemiopsis leidyi* MA  *This study* | *Mnemiopsis gardeni* FL  *This study* | *Mnemiopsis leidyi*  Germany  Koutsouveli et al. 2025 |
| --- | --- | --- | --- | --- |
| Assembly size | 155.8 Mb | 215.8 Mb | 206.6 Mb | 224.5 Mb |
| No. of scaffolds | 5,100 | NA | NA | 13 |
| No. of contigs | NA | 207 | 174 | NA |
| N50 | 187 Kb (scaff) | 2 Mb (contig) | 3.6 Mb (contig) | 17.3 Mb (scaff) |
| BUSCO Complete | 204 (80%) | 211 (83%) | 214 (84%) | 208 (82%) |
| BUSCO Complete + Partial | 233 (91%) | 231 (91 %) | 235 (92%) | 230 (90%) |
| BUSCO missing | 22 (8.6%) | 24 (9%) | 20 (8%) | 25 (10%) |
| % of detected core genes with > 1 ortholog | 1.47 | 1.9 | 3.74 | 4.33 |

**Supplemental Table 3.** Assembled gene/protein models that were annotated using three protein databases: UniProt Knowledgebase Swiss-Prot protein models v2021-03, RefSeq invertebrate protein models, and the *M. leidyi* protein models.

**Supplemental Table 4.** F_ST_ values across all populations of *Mnemiopsis* collected based on the LD-Filtered SNP dataset. These locations included: Woods Hole, MA (MAW), Little Egg Harbor Township, NJ (NJR), Berlin, MD (MDA), Gloucester Point, VA (VAV), Roanoke Island, NC (NCX), Pivers Island, NC (NCB), Wrightsville Beach, NC (NCW), Charleston, SC (SCC), Savannah, GA (GAS), Savannah offshore, GA (GAB), Flagler Beach, FL (FLM), Fort Pierce, FL (FLF), and Panacea, FL (FLP).

|  | MAW | NJR | MDA | VAV | NCX | NCB | NCW | SCC | GAS | GAB | FLM | FLF | FLP |
| --- | --- | --- | --- | --- | --- | --- | --- | --- | --- | --- | --- | --- | --- |
| MAW |  |  |  |  |  |  |  |  |  |  |  |  |  |
| NJR | 0.039 |  |  |  |  |  |  |  |  |  |  |  |  |
| MDA | 0.056 | -0.003 |  |  |  |  |  |  |  |  |  |  |  |
| VAV | 0.215 | 0.097 | 0.075 |  |  |  |  |  |  |  |  |  |  |
| NCX | 0.327 | 0.230 | 0.210 | 0.113 |  |  |  |  |  |  |  |  |  |
| NCB | 0.400 | 0.328 | 0.314 | 0.262 | 0.230 |  |  |  |  |  |  |  |  |
| NCW | 0.402 | 0.330 | 0.317 | 0.266 | 0.235 | 0.001 |  |  |  |  |  |  |  |
| SCC | 0.402 | 0.331 | 0.318 | 0.266 | 0.236 | 0.0003 | 0.0002 |  |  |  |  |  |  |
| GAS | 0.407 | 0.335 | 0.322 | 0.271 | 0.240 | 0.001 | 0.001 | 0.001 |  |  |  |  |  |
| GAB | 0.402 | 0.330 | 0.317 | 0.265 | 0.235 | -0.0003 | 0.001 | 0.00004 | −0.00002 |  |  |  |  |
| FLM | 0.391 | 0.318 | 0.305 | 0.253 | 0.223 | 0.029 | 0.028 | 0.027 | 0.032 | 0.027 |  |  |  |
| FLF | 0.404 | 0.331 | 0.318 | 0.265 | 0.235 | 0.061 | 0.058 | 0.087 | 0.064 | 0.057 | 0.03 |  |  |
| FLP | 0.422 | 0.349 | 0.336 | 0.283 | 0.254 | 0.089 | 0.086 | 0.087 | 0.092 | 0.086 | 0.059 | 0.039 |  |

**Supplementary Table 5.** *f*_3_ results where each row represents a scenario where the FLP and MAW populations are the source for admixture and the third population is the target of hybridization. A *f*_3_ statistic, the standard error (SE) and a Z-score value calculated with jackknife is provided for each scenario.

|  | A | B | C | *f*_3_ | stderr | Zscore | Nsnps |
| --- | --- | --- | --- | --- | --- | --- | --- |
| 1 | FLP | MAW | NJR | -0.0737 | 0.00264 | -28.0 | 4353101 |
| 2 | FLP | MAW | MDA | -0.0842 | 0.00281 | -30.0 | 4366064 |
| 3 | FLP | MAW | VAV | -0.0424 | 0.00335 | -12.7 | 4420740 |
| 4 | FLP | MAW | NCX | -0.0233 | 0.00589 | -3.96 | 4457812 |
| 5 | FLP | MAW | NCB | 0.0406 | 0.00795 | 5.11 | 4711795 |
| 6 | FLP | MAW | NCW | 0.0464 | 0.00754 | 6.16 | 4710393 |
| 7 | FLP | MAW | SCC | 0.0460 | 0.00723 | 6.36 | 4709074 |
| 8 | FLP | MAW | GAS | 0.0542 | 0.00881 | 6.16 | 4720780 |
| 9 | FLP | MAW | GAB | 0.0455 | 0.00738 | 6.16 | 4710162 |
| 10 | FLP | MAW | FLM | 0.0142 | 0.00260 | 5.47 | 4676756 |
| 11 | FLP | MAW | FLF | 0.00534 | 0.00125 | 4.27 | 4584377 |

**Supplemental Table 6.** The top 1% of fixation index values (*V*_ST_) for the copy number variation analyses with their associated annotations taken from Supplemental Table 3.

**Supplemental Table 7.** Gene Ontology (GO) analysis of genes in the top 1% regions of copy number variation outliers reveals enrichment in molecular functions (MF), cellular components (CC), and biological processes (BP).

| *Rank* | *GO ID* | *Category* | *Num in category* | *P-value* | *GO Description* |
| --- | --- | --- | --- | --- | --- |
| 1 | GO:0004819 | MF | 1 | 0.0023 | glutamine-tRNA ligase activity |
| 2 | GO:0006425 | BP | 1 | 0.0023 | glutaminyl-tRNA aminoacylation |
| 3 | GO:0015074 | BP | 3 | 0.0113 | DNA integration |
| 4 | GO:0016876 | MF | 1 | 0.0241 | aminoacyl-tRNA ligase activity |
| 5 | GO:0043039 | BP | 1 | 0.0345 | tRNA aminoacylation |
| 6 | GO:0003676 | MF | 7 | 0.0357 | nucleic acid binding |
| 7 | GO:0007155 | BP | 1 | 0.0600 | cell adhesion |
| 8 | GO:0004523 | MF | 1 | 0.0670 | RNA-DNA hybrid ribonuclease activity |
| 9 | GO:0016779 | MF | 1 | 0.0725 | nucleotidyltransferase activity |
| 10 | GO:0006418 | BP | 1 | 0.1074 | tRNA aminoacylation for protein translation |
| 11 | GO:0004812 | MF | 1 | 0.1170 | aminoacyl-tRNA ligase activity |
| 12 | GO:0046983 | MF | 1 | 0.2033 | protein dimerization activity |
| 13 | GO:0005509 | MF | 2 | 0.2112 | calcium ion binding |
| 14 | GO:0000166 | MF | 1 | 0.2414 | nucleotide binding |
| 15 | GO:0006412 | BP | 1 | 0.3122 | translation |
| 16 | GO:0008270 | MF | 2 | 0.3299 | zinc ion binding |
| 17 | GO:0004930 | MF | 3 | 0.4098 | G protein-coupled receptor activity |
| 18 | GO:0007186 | BP | 3 | 0.4221 | G protein-coupled receptor signaling pathway |
| 19 | GO:0005622 | CC | 1 | 0.4512 | intracellular anatomical structure |
| 20 | GO:0005737 | CC | 1 | 0.4647 | cytoplasm |
| 21 | GO:0046872 | MF | 1 | 0.4713 | metal ion binding |
| 22 | GO:0016021 | CC | 5 | 0.5852 | membrane |
| 23 | GO:0055085 | BP | 1 | 0.7341 | transmembrane transport |
| 24 | GO:0016491 | MF | 1 | 0.8124 | oxidoreductase activity |
| 25 | GO:0005515 | MF | 4 | 0.8238 | protein binding |
| 26 | GO:0055114 | BP | 1 | 0.8908 | obsolete oxidation-reduction process |
| 27 | GO:0005524 | MF | 1 | 0.9378 | ATP binding |

**Supplemental Table 8.** Putative G protein-coupled receptors in the top 1% of copy number variation outliers.

| *ID* | *BEST UNIPROT HIT* |
| --- | --- |
| jg9426 | BRS3_SHEEP |
| jg3742 | CLTR2_RAT |
| jg17847 | GPR19_MOUSE |
| jg14707 | QRFPR_BRAFL |

**Supplemental Table 9.** The top 10% of XP-CLR values calculated between the two lineages using a sliding window approach, a step-size of 10 Kb and the maxSNPs flag set to 200. We have appended their associated annotations from Supplemental Table 3.

**Supplemental Table 10.** Putative G-coupled protein receptors in the top 10% of XP-CLR outliers with corresponding *M. gardeni* gene ID.

| *ID* | *BEST UNIPROT HIT* |
| --- | --- |
| jg24766 | OPSD_SPHSP |
| jg17906 | OPSP_COLLI |
| jg18796 | MTR1B_RAT |
| jg22094 | ADA1A_ORYLA |
| jg9318 | CLTR2_MOUSE |
| jg8157 | NPR22_CAEEL |
| jg9346 | OPSD1_MIZYE |
| jg17110 | OPSD1_MIZYE |
| jg15114 | GR101_LYMST |
| jg4357 | CXCR1_RAT |
| jg3032 | GR101_LYMST |
| jg12031 | 5HT1A_MOUSE |
| jg16938 | GR101_LYMST |
| jg6892 | GR101_LYMST |
| jg21723 | CLTR1_RAT |
| jg9722 | C5AR1_CAVPO |
| jg10869 | GAL2B_DANRE |
| jg4732 | GPR17_HUMAN |
| jg11407 | QRFPR_BRAFL |
| jg5542 | TLR1_DROME |
| jg6417 | CLTR1_RAT |
| jg6367 | OPSD1_MIZYE |
| jg3602 | GR101_LYMST |
| jg24926 | OX2R_MOUSE |
| jg18531 | GR101_LYMST |
| jg7508 | ADRB1_MELGA |
| jg15516 | ADA2B_DANRE |
| jg15089 | FFAR3_RAT |
| jg6968 | CXCR2_RAT |
| jg13039 | OR7C2_HUMAN |
| jg9196 | HRH2_PONPY |
| jg3259 | PRLHR_MOUSE |
| jg21687 | OPN4_FELCA |
| jg20054 | GR101_LYMST |
| jg20383 | PAR2_MOUSE |
| jg12315 | AGRD1_BOVIN |
| jg11079 | GP83A_DANRE |
| jg20519 | GR101_LYMST |
| jg17168 | OPSD1_MIZYE |
| jg16790 | ADA1A_CAVPO |
| jg4268 | GPR17_RAT |
| jg8834 | GPR87_HUMAN |
| jg21572 | GPR25_HUMAN |
| jg21573 | P2RY4_HUMAN |
| jg14193 | CLTR1_CAVPO |
| jg15046 | ADA1B_RAT |
| jg8705 | ADB4C_MELGA |
| jg10820 | NPFR_DROME |
| jg21670 | CCH1R_DROME |
| jg23090 | GAL2B_DANRE |
| jg23091 | CLTR1_CAVPO |
| jg7663 | GPR21_HUMAN |
| jg23495 | GP183_HUMAN |
| jg20840 | GR101_LYMST |
| jg23465 | GRM7_PONAB |
| jg21077 | CXCR1_RABIT |
| jg10716 | CXR4A_XENLA |
| jg23083 | GP183_RAT |
| jg11802 | OPSD_ALLMI |
| jg21919 | OPSD_ASTFA |
| jg8302 | OPN4_HUMAN |
| jg4304 | P2RY1_CHICK |
| jg21417 | MTR1L_HUMAN |
| jg20841 | P2RY1_HUMAN |
| jg18789 | CCR4_MOUSE |
| jg23277 | OPRK_RAT |
| jg7942 | OPSD1_MIZYE |
| jg2813 | GR101_LYMST |
| jg24911 | OPSD1_MIZYE |
| jg9218 | SIFAR_DROME |
| jg23329 | CCKAR_MOUSE |
| jg19814 | GR101_LYMST |
| jg8431 | GALR2_HUMAN |
| jg11123 | GP161_BOVIN |
| jg2735 | GRIK2_MACFA |
| jg7993 | HRH2_MOUSE |
| jg7407 | AGRD1_BOVIN |
| jg4339 | GRM3_PONAB |
| jg7991 | S39AB_HUMAN |
| jg13104 | MTR1C_XENLA |
| jg15080 | GR101_LYMST |
| jg21725 | GPC6A_DANRE |
| jg5285 | GR101_LYMST |
| jg24843 | NPFF2_HUMAN |
| jg21456 | GPC6A_MOUSE |
| jg5225 | HRH2_MOUSE |
| jg20562 | GR101_LYMST |
| jg14575 | CCR4_HUMAN |
| jg23425 | LRP2_RAT |
| jg20230 | OPSD1_MIZYE |
| jg17099 | OPRK_CAVPO |
| jg3722 | EGG1_CAERE |
| jg12468 | ADA1D_HUMAN |

**Supplemental Table 11.** Channels and receptors in the top 10% of XP-CLR outliers with corresponding *M. gardeni* gene ID.

| *ID* | *Type* | *Top UniProt hit* |
| --- | --- | --- |
| jg12907 | 2-pore calcium channel | TPC1_ARATH |
| jg9066 | Acid-sensing ion channels | ASIC4_RAT |
| jg9132 | Acid-sensing ion channels | ASI4A_DANRE |
| jg9383 | Acid-sensing ion channels | ASIC5_MOUSE |
| jg7342 | Calcium-activated K+ channel | KCMA1_BOVIN |
| jg9510 | Calcium-activated K+ channel | KCMB3_HUMAN |
| jg11518 | DEG/ENaC channel | DEL1_CAEEL |
| jg11565 | Glutamate receptor | GRIK2_HUMAN |
| jg2735 | Glutamate receptor | GRIK2_MACFA |
| jg3929 | Glutamate receptor | GRIK2_XENLA |
| jg6697 | Glutamate receptor | GRIK4_PANTR |
| jg8199 | Glutamate receptor | GRIK2_MOUSE |
| jg3342 | Synaptojanin | SYJ2B_RAT |
| jg7623 | Syntaxin | STX6_RAT |
| jg22067 | TRPC | TRPC4_HUMAN |
| jg22067 | TRPC | TRPC4_HUMAN |
| jg22066 | TRPL | TRPL_CAEEL |
| jg22457 | TRPM | TRPM2_HUMAN |
| jg18519 | TWiK potassium channel | TWK7_CAEEL |
| jg11558 | Voltage-gated chloride channel | CLCN2_CAVPO |
| jg5899 | Voltage-gated sodium channel | SC4AB_DANRE |
| jg23380 | Voltage-gated calcium channel | CAC1D_DROME |
| jg11179 | Voltage-gated potassium channel | KCNA1_MOUSE |
| jg11559 | Voltage-gated potassium channel | KCNA2_HUMAN |
| jg15226 | Voltage-gated potassium channel | KCNA1_MOUSE |
| jg19404 | Voltage-gated potassium channel | KCNA5_HUMAN |
| jg19411 | Voltage-gated potassium channel | KCNA1_HUMAN |
| jg20392 | Voltage-gated potassium channel | KCNA2_ONCMY |
| jg20686 | Voltage-gated potassium channel | KCNA3_HUMAN |
| jg20950 | Voltage-gated potassium channel | KCNA1_HUMAN |
| jg21641 | Voltage-gated potassium channel | KCNA2_ONCMY |

**Supplemental Table 12.** Gene Ontology (GO) analysis of genes in the top 10% of XP-CLR outlier regions reveals enrichment in molecular functions, cellular components, and biological processes.

| *Rank* | *GO ID* | *Category* | *Num in category* | *P-value* | *GO Description* |
| --- | --- | --- | --- | --- | --- |
| 1 | GO:0005515 | MF | 291 | 0.0000 | protein binding |
| 2 | GO:0003723 | MF | 43 | 0.0000 | RNA binding |
| 3 | GO:0030117 | CC | 6 | 0.0005 | membrane coat |
| 4 | GO:0004672 | MF | 51 | 0.0007 | protein kinase activity |
| 5 | GO:0006468 | BP | 51 | 0.0012 | protein phosphorylation |
| 6 | GO:0005488 | MF | 42 | 0.0019 | binding |
| 7 | GO:0015035 | MF | 10 | 0.0029 | protein-disulfide reductase activity |
| 8 | GO:0007018 | BP | 12 | 0.0048 | microtubule-based movement |
| 9 | GO:0006662 | BP | 4 | 0.0054 | glycerol ether metabolic process |
| 10 | GO:0008061 | MF | 9 | 0.0056 | chitin binding |
| 11 | GO:0045454 | BP | 17 | 0.0066 | cell redox homeostasis |
| 12 | GO:0003755 | MF | 6 | 0.0066 | peptidyl-prolyl cis-trans isomerase activity |
| 13 | GO:0000413 | BP | 6 | 0.0066 | protein peptidyl-prolyl isomerization |
| 14 | GO:0006379 | BP | 3 | 0.0069 | obsolete mRNA cleavage |
| 15 | GO:0006452 | BP | 2 | 0.0069 | translational frameshifting |
| 16 | GO:0045905 | BP | 2 | 0.0069 | positive regulation of translational termination |
| 17 | GO:0045901 | BP | 2 | 0.0069 | positive regulation of translational elongation |
| 18 | GO:0006030 | BP | 7 | 0.0073 | chitin metabolic process |
| 19 | GO:0005216 | MF | 21 | 0.0080 | monoatomic ion channel activity |
| 20 | GO:0005868 | CC | 2 | 0.0087 | cytoplasmic dynein complex |
| 21 | GO:0044877 | MF | 2 | 0.0090 | protein-containing complex binding |
| 22 | GO:0006811 | BP | 21 | 0.0095 | monoatomic ion transport |
| 23 | GO:0042393 | MF | 2 | 0.0104 | histone binding |
| 24 | GO:0006886 | BP | 13 | 0.0105 | intracellular protein transport |
| 25 | GO:0006836 | BP | 5 | 0.0121 | neurotransmitter transport |
| 26 | GO:0043022 | MF | 3 | 0.0135 | ribosome binding |
| 27 | GO:0007186 | BP | 102 | 0.0136 | G protein-coupled receptor signaling pathway |
| 28 | GO:0004930 | MF | 100 | 0.0152 | G protein-coupled receptor activity |
| 29 | GO:0006820 | BP | 4 | 0.0152 | monoatomic anion transport |
| 30 | GO:0030246 | MF | 11 | 0.0154 | carbohydrate binding |
| 31 | GO:0006396 | BP | 10 | 0.0176 | RNA processing |
| 32 | GO:0016021 | CC | 205 | 0.0178 | membrane |
| 33 | GO:0003676 | MF | 132 | 0.0183 | nucleic acid binding |
| 34 | GO:0031072 | MF | 2 | 0.0232 | heat shock protein binding |
| 35 | GO:0008290 | CC | 2 | 0.0243 | F-actin capping protein complex |
| 36 | GO:0051016 | BP | 2 | 0.0243 | barbed-end actin filament capping |
| 37 | GO:0006270 | BP | 4 | 0.0248 | DNA replication initiation |
| 38 | GO:0006383 | BP | 2 | 0.0250 | transcription by RNA polymerase III |
| 39 | GO:0006813 | BP | 12 | 0.0278 | potassium ion transport |
| 40 | GO:0042555 | CC | 3 | 0.0284 | MCM complex |
| 41 | GO:0003777 | MF | 10 | 0.0308 | microtubule motor activity |
| 42 | GO:0051056 | BP | 3 | 0.0322 | regulation of small GTPase mediated signal transduction |
| 43 | GO:0008076 | CC | 10 | 0.0334 | voltage-gated potassium channel complex |
| 44 | GO:0035556 | BP | 14 | 0.0355 | intracellular signal transduction |
| 45 | GO:0005622 | CC | 29 | 0.0398 | intracellular anatomical structure |
| 46 | GO:0048193 | BP | 2 | 0.0399 | Golgi vesicle transport |
| 47 | GO:0004435 | MF | 2 | 0.0408 | phosphatidylinositol phospholipase C activity |
| 48 | GO:0008081 | MF | 7 | 0.0420 | phosphoric diester hydrolase activity |
| 49 | GO:0005328 | MF | 4 | 0.0424 | neurotransmitter:sodium symporter activity |
| 50 | GO:0005249 | MF | 10 | 0.0443 | voltage-gated potassium channel activity |
| 51 | GO:0007205 | BP | 2 | 0.0444 | protein kinase C-activating G protein-coupled receptor  signaling pathway |
| 52 | GO:0004143 | MF | 2 | 0.0444 | ATP-dependent diacylglycerol kinase activity |
| 53 | GO:0009055 | MF | 10 | 0.0454 | electron transfer activity |
| 54 | GO:0006108 | BP | 2 | 0.0457 | malate metabolic process |
| 55 | GO:0016615 | MF | 2 | 0.0457 | malate dehydrogenase activity |
| 56 | GO:0031177 | MF | 2 | 0.0460 | phosphopantetheine binding |
| 57 | GO:0008509 | MF | 2 | 0.0475 | monoatomic anion transmembrane transporter activity |
| 58 | GO:0008017 | MF | 8 | 0.0486 | microtubule binding |

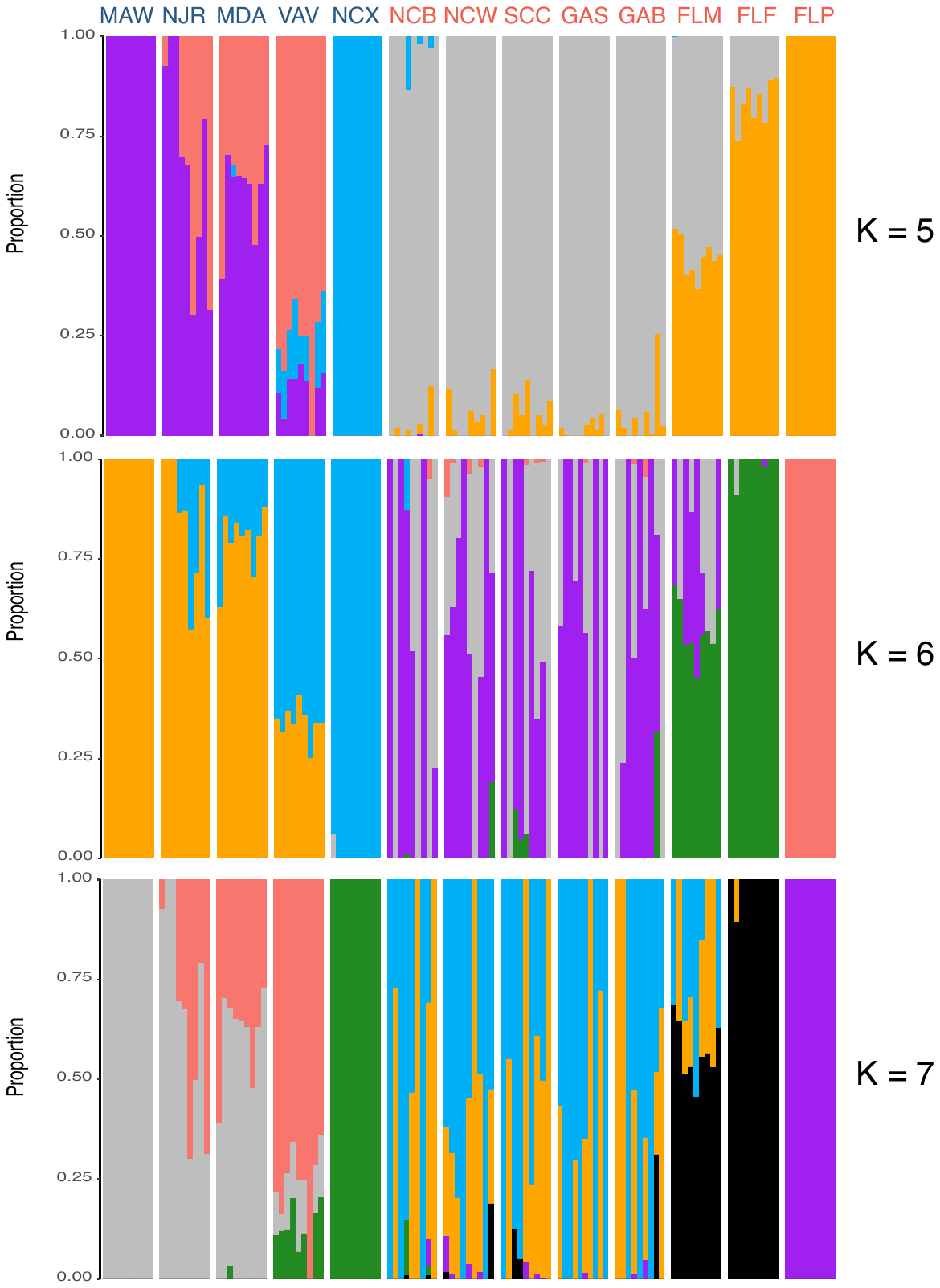

**Supplemental Figure 1.** Admixture plots for the 118 *Mnemiopsis* samples collected for K = 5 to K = 7 where each bar represents a single individual, and the colors represent each of the K components.

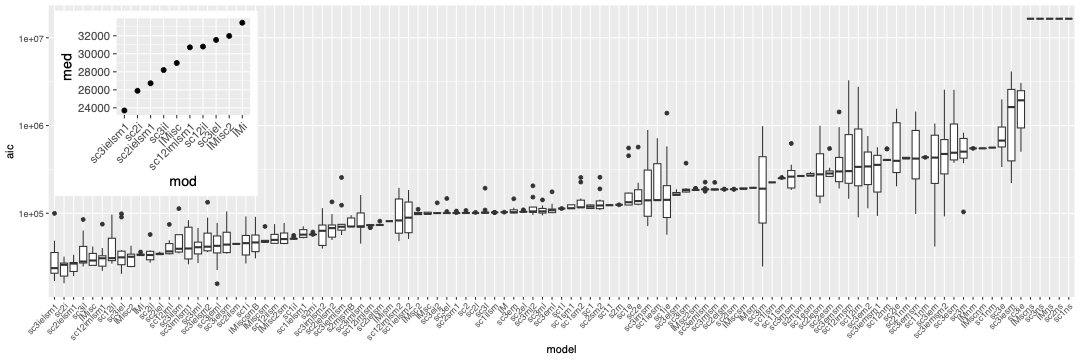

**Supplemental Figure 2.** Boxplots summarizing the best AIC scores for each model for all bootstrapped replicates for each of 102 demographic models tested. The inset plot shows the boxplot for the median AIC for the top ten best-fitting models in the set.

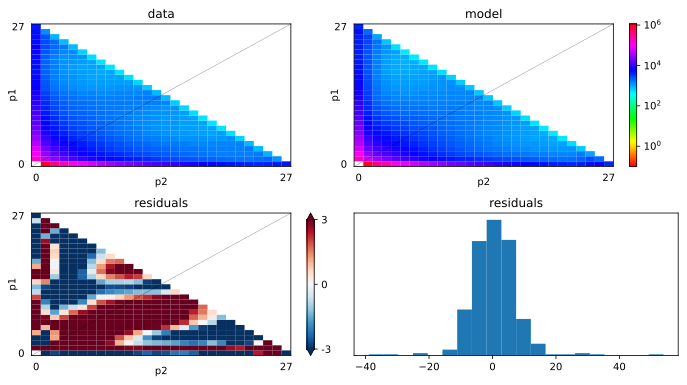
**Supplemental Figure 3.** Plots showing the goodness-of-fit of the best fitting model (sc3ielsm1). The top two panels show the observed AFS (top left) and the modeled AFS (top right). The lower two plots show model errors plotted as residuals between the best-fit model and the data (bottom left) and the frequency histogram of residuals (bottom right).

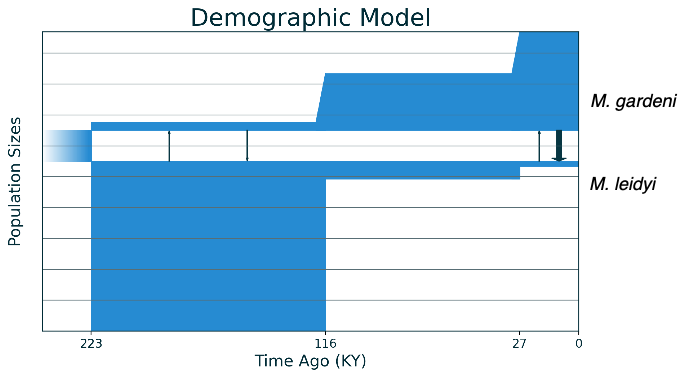

**Supplemental Figure 4.** The best-fit demographic model (sc3ielsm1) inferred using *Moments* for the demographic history between *M. leidyi* and *M. gardeni*. This best-fit model includes three epochs in each species with symmetric migration in the first and asymmetric migration in the last.

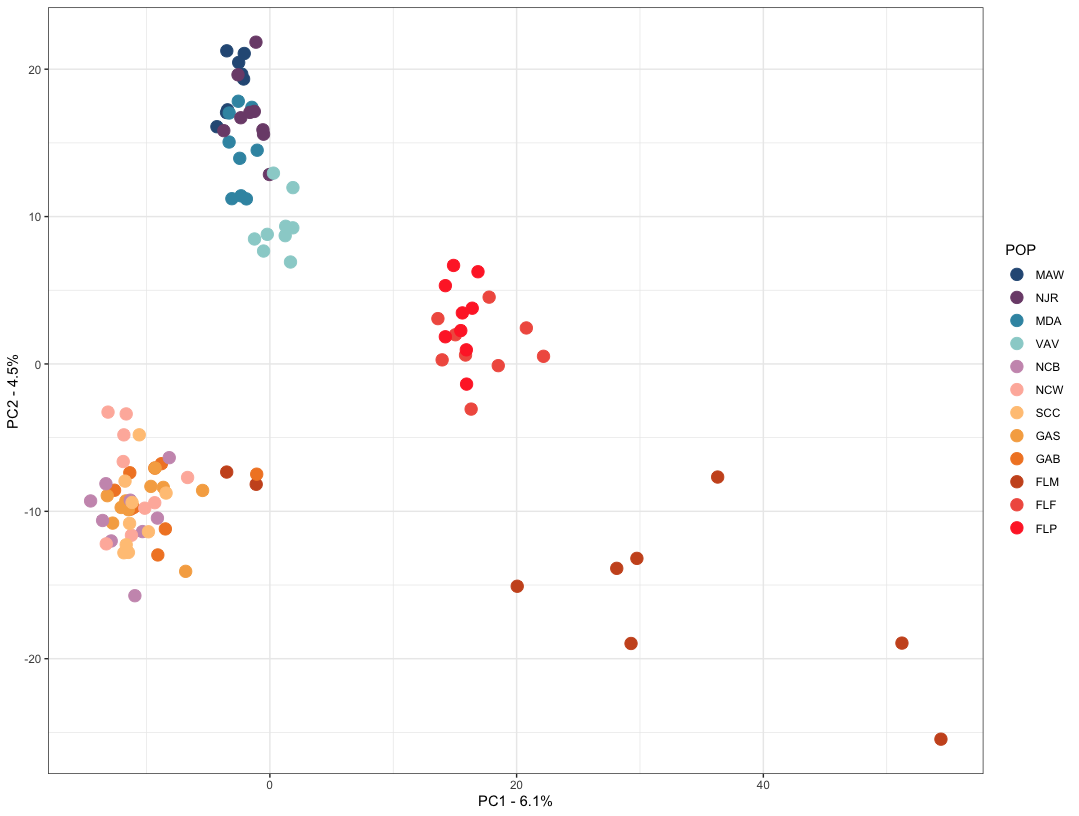
**Supplemental Figure 5.** PCA plot based on copy number variation among samples. Copy number estimates were generated as denoised copy number ratios in GATK. Individuals are colored by their sampling location.

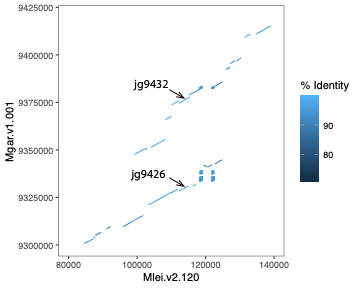

**Supplemental Figure 6.** The top *V*_ST_ outlier gene occurs in an *M. gardeni*-specific duplicated region. This dot plot shows BLAST hits (e-value < 1e-10; length >= 500bp) between corresponding regions in the *M. leidyi* and *M. gardeni* assemblies. The *V*_ST_ outlier gene (jg9426 – Mgar.v1.001:9329910-9331046) occurs as part of a larger duplicated region in *M. gardeni*. In *M. leidyi,* this region is single-copy. Arrows highlight the location of the *M. gardeni* duplicated genes.
